# *In situ* generation of RNA complexes for synthetic molecular strand displacement circuits in autonomous systems

**DOI:** 10.1101/2020.07.15.204438

**Authors:** Wooli Bae, Guy-Bart V. Stan, Thomas E. Ouldridge

**Author notes:** Correspondence and request for materials should be addressed to G.-B.S. and T.O.

## Abstract

Synthetic molecular circuits implementing DNA or RNA strand-displacement reactions can be used to build complex systems such as molecular computers and feedback control systems. Despite recent advances, application of nucleic acid-based circuits *in vivo* remains challenging due to a lack of efficient methods to produce their essential components – multi-stranded complexes known as “gates” – *in situ*, i.e. in living cells or other autonomous systems. Here, we propose the use of naturally occurring self-cleaving ribozymes to cut a single-stranded RNA transcript into a gate complex of shorter strands, thereby opening new possibilities for the autonomous and continuous production of RNA strands in a stoichiometrically and structurally controlled way.

---

Nucleic acid molecules offer a unique feature that make them ideal for designing nanostructures and synthetic molecular reactions: predictable Watson-Crick base pairing interactions^1, 2^. By designing oligonucleotide molecules with specific sequences, it is possible to build complex synthetic systems where the domains of designed nucleotide sequences interact in a predictable and programmable manner^3-5^. One of the core reactions in such synthetic nucleic acid circuits is toehold-mediated strand-displacement^6-8^. In a typical toehold-mediated strand displacement reaction, an input strand binds to a pre-formed multi-stranded complex via a toehold region. Binding of the input triggers the displacement of an output strand, which can then serve as an input strand for other reactions. This programmable input-output process can be abstracted to a functional logic gate with the multi-stranded complex being a “gate”. With sets of engineered toehold-mediated strand-displacement reactions using the gates, computationally generalizable and composable molecular reaction networks of increasing complexity have been proposed^9^, incorporating multi-input, multi-output gates and components that act as molecular machines^10, 11^ and perform complex computations^12-16^. In such reaction networks, the engineered nucleic acid sequences not only dictate the order and rate of the reactions but also drive these reactions forward via the lower free energy of the reaction products.

Motivated by the potential of nucleic-acid-based systems for predictable and programmable behaviours, research has focused on their use in synthetic biology to engineer and control molecular systems implemented in living organisms^17, 18^. Much of the work on implementing nucleic acid nanotechnology in living cells has so far focussed on the assembly^19-21^ or delivery of static structures^22, 23^. Dynamic circuitry has hitherto either not used strand displacement^24^, relied on exogenously supplied gates^25^ or used displacement only to modify secondary structure in single strands^26-29^ Although powerful, these formalisms lack the composability and simplicity of reaction networks based on strand displacement gates. Exogenously supplied gates offer such composability and simplicity but suffer from low delivery efficiency and toxicity from the delivering agents^30^. Moreover, external delivery of components is not a sustainable strategy for the autonomous, long-term operation of circuits, especially for a system with large number of cells. Therefore, there is a clear need for a method that allows for the autonomous production of gates.

The current methodology used to produce strand displacement-ready gates relies on chemical synthesis of individual oligonucleotides followed by mixing and thermal annealing (Figure 1, upper panel). Unfortunately, this approach is incompatible with the production of gates in autonomous systems such as living cells. Furthermore, the naive strategy of expressing *individual* strands via transcription, and relying on them to form gate complexes *in situ*, is flawed. First, a basic premise of strand displacement circuits is that thermodynamically favourable reactions are kinetically supressed by the sequestration of strands in gate complexes; these reactions only occur fast if appropriate triggers are present. If strands were transcribed individually and simultaneously, there would be nothing to prevent the direct formation of products in the absence of a trigger, unless the initial components were mixed in separate compartments^31^. Secondly, it is difficult to achieve precise stoichiometry of the various strands generated from different pathways, which is typically required to prevent leak reactions^32^.

**Figure 1.**
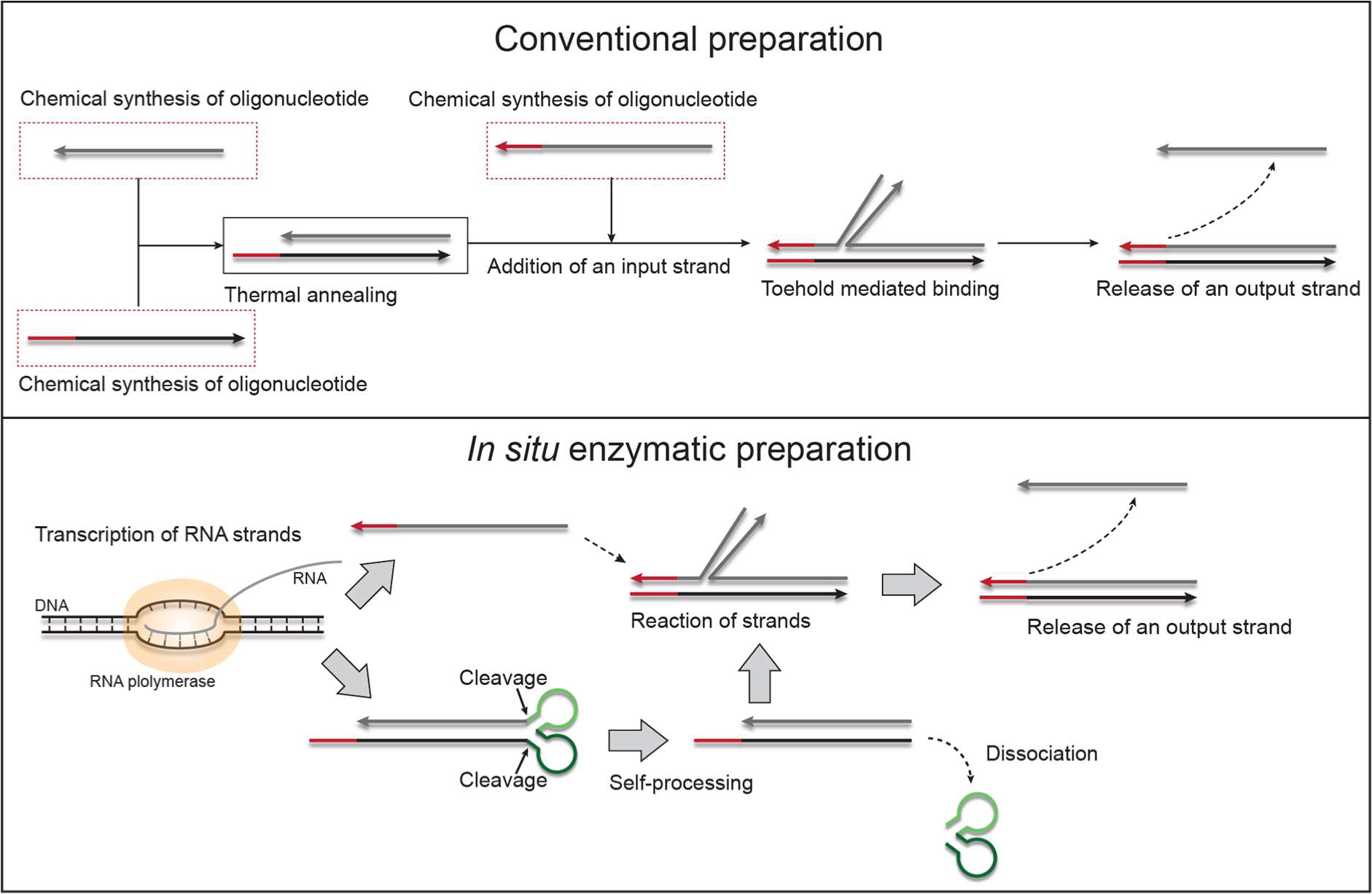
Comparison of conventional preparation methods with the *in situ* enzymatic preparation of multi-stranded RNA oligonucleotides. With the conventional method (upper panel), each oligonucleotide is synthesized chemically and then thermally annealed in a separate process to form a multi-stranded complex. Another chemically synthesized input strand is then added to trigger a toehold-mediated strand-displacement reaction that releases an output strand. Each rectangular box represents a separate reaction pot. With our *in situ* enzymatic method (lower panel), every RNA strand is enzymatically synthesized from its corresponding DNA template in the same volume. The RNAs fold into designed secondary structures harbouring self-cleaving hammerhead ribozymes (HHR, green domains). After self-cleavage upstream near the 5’ end (light green domain) and downstream near the 3’ end (dark green domain) of the HHR domain, the self-excising module dissociates from the rest of the RNA leaving a multi-stranded RNA. This multi-stranded complex reacts with an input strand to release an output strand.

Here, we propose direct production of ribonucleotide gates in an *in vitro* transcription system by transcribing a long RNA strand that cuts itself into separate strands via the action of self-cleaving ribozymes^33, 34^. A pair of ribozymes are designed to form a self-excising module that cuts itself out from the rest of the gates. We first demonstrate that the self-excising module successfully cleaves itself and dissociates from the rest of the RNA transcript, leaving only the gate. We then demonstrate the *in situ* production and operation of a single-input, single-output gate and its trigger molecule, and the orthogonality of two trigger/gate pairs. Finally, we demonstrate *in situ* production and operation of a two-step cascade circuit where one gate is triggered by the output of another gate.

In our *in situ* enzymatic preparation scheme (Figure 1, lower panel), an RNA strand is transcribed from a DNA template that encodes the sequence for single-stranded inputs or gate complexes. For the gates, the transcribed RNA sequences are designed to fold into specific gate-like secondary structures, but with the strand ends connected by a self-excising module consisting of two ribozymes. Each self-excising module is designed to cut the RNA backbone twice and dissociate completely from the rest of the gate. With their ends disconnected, the gate complexes can be used in normal strand-displacement reactions. All the reactions, including synthesis, folding and self-cleavage of RNA happen autonomously, i.e. without any external input.

The self-excising module was designed by directly connecting two *cis*-acting HHRs that cleave upstream near the 5’ ends (HHR1, from chrysanthemum chlorotic mottle viroid^33^) or downstream near the 3’ ends (HHR2, from Schistosoma mansoni^34^, Figure 2a). We first tested the activity of each HHR separately. The corresponding DNA templates were first constructed and amplified (Figure S1). After two hours of transcription, each RNA was run in a denaturing PAGE gel to check its cleavage yield (Figures 2b, c). In both cases, we observed successful cleavage as indicated by the appearance of additional PAGE gel bands relative to the control consisting of a deactivated (Figure S2) ribozyme.

**Figure 2.**
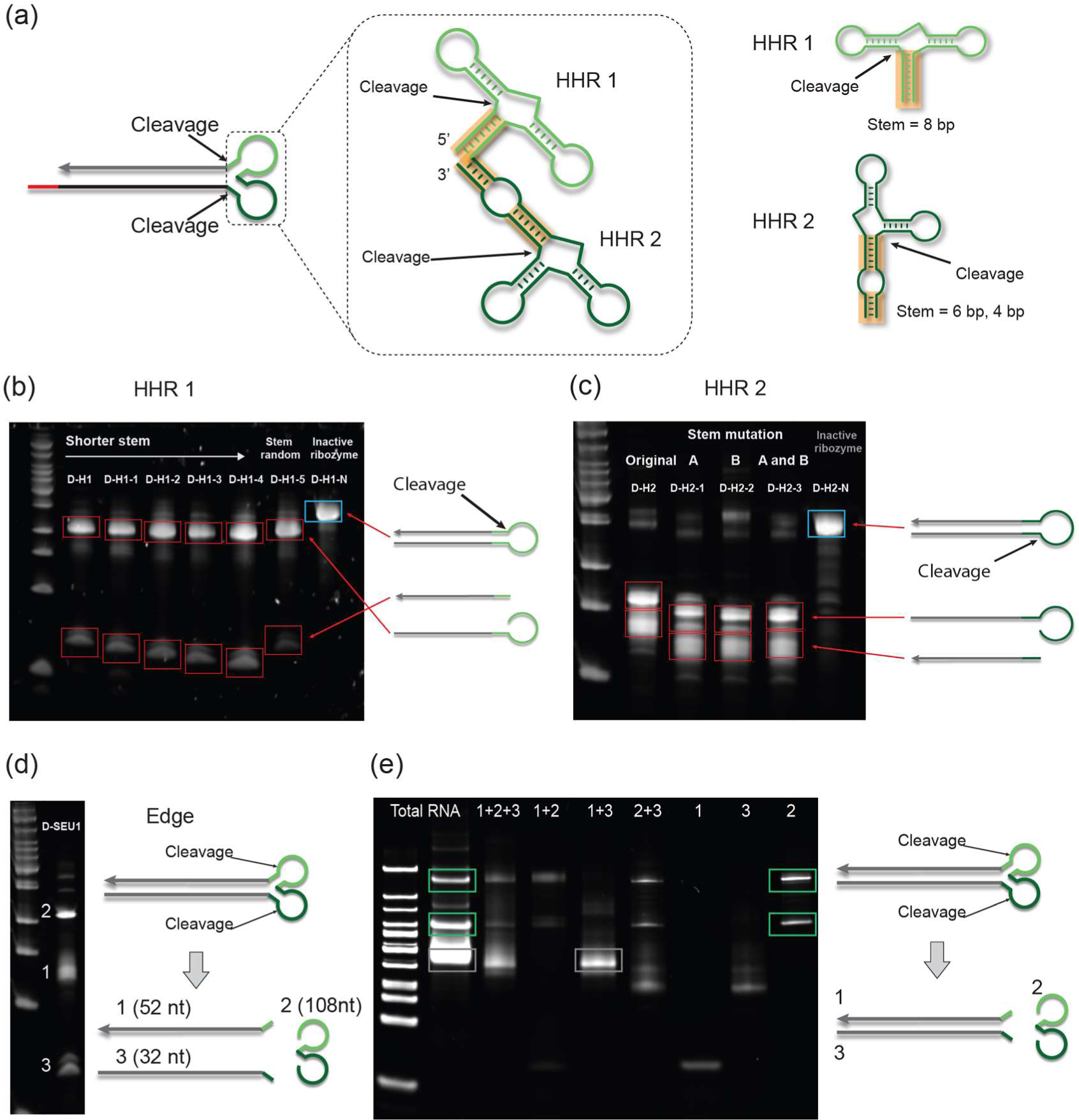
Engineering hammerhead ribozyme (HHR) components for the self-excising ribozyme module. (a) The self-excising module consists of two HHRs that cleave themselves near the 5’ (HHR1, from *chrysanthemum chlorotic mottle viroid* ^*33*^) and 3’ (HHR2, from *Schistosoma mansoni* ^*34*^) ends, respectively. (b, c) The activity of each ribozyme was checked by running the transcribed RNA from each DNA template into a denaturing PAGE gel. A successful cleavage reaction is marked by the appearance of an additional band and a shift of the main band (red squares) compared to a transcript with a non-functional ribozyme (blue squares). Lanes are labelled by the DNA construct (sequences in supplementary Excel file), constructs with shortened (7-4 bp) or randomized stems showed similar cleavage yield. (d) The self-excising module was tested similarly. Successful cleavage was observed through the appearance of three major bands that correspond to their size. (e) The RNA transcript from the self-excising module template (D-SEU1)was run on a native PAGE gel alongside mixtures of separately produced control RNA transcripts to test whether the self-excising module spontaneously dissociates from the rest of the transcript after cleavage. The bands observed for full RNA transcripts correspond to the superposition of those obtained from a self-excising HHR (green boxes) and a multi-stranded RNA complex (grey boxes), showing that the self-excising HHR will spontaneously dissociate from a duplex in non-denaturing conditions. Full description of the DNA templates used in the figure can be found in the Supplementary Excel file.

Based on the relative intensity of these bands, we estimated the cleavage yield of each HHR to be roughly 90%, similar to the previously reported yield for full-length HHRs (see Supplementary methods and Supplementary data). We then tested if we could modify the sequence of the stem (orange boxes in Figure 2a) as this region is retained as a part of the multi-stranded complex after dissociation of the self-excising module. We reasoned that the stem region is just a scaffold that stabilizes the tertiary structure of the HHR since the sequence of the stem is not conserved among different species of HHRs^35^. Indeed, shortening the stem of HHR1 by up to 4 base pairs or replacing the sequence of the stem with a random sequence did not hamper cleavage (Figure 2b). As the HHR2 has two shorter stems (6 bp and 4 bp long, respectively) with a functional loop in between, instead of one long stem, we tried replacing the sequence of either or both stems (Figure 2a, orange boxes in dark green structure). This did not hamper cleavage (Figure 2c).

After confirming the activity of each HHR, we assembled the self-excising module by directly connecting HHR2 to the 3’ end of HHR1 (Figure 2a). With two ribozymes on the RNA transcript, two successful cleavage events on the self-excising module result in three separate strands, which migrate as three major bands in a denaturing gel (Figure 2d). The estimated yield for a complete cleavage reaction of the self-excising module was roughly 80% (Figure S3), which is lower than the cleavage yield obtained from a single HHR. Since this yield is simply a combination yield for two independent cleavage processes with ∼90% yield for each, the activity of each HHR is not substantially reduced by connecting them together. Furthermore, shortening the stem of HHR1 or replacing the sequence of the stems of each HHR did not substantially reduce the cleavage yield of the module (Figure S3). Even after successful cleavage reactions, however, it is possible that the self-excising module remains bound to the rest of the RNA complex using residual base-pairing interactions in non-denaturing conditions (Figure 2a, orange boxes). To check this behaviour, we ran the RNA transcript harbouring a self-excising module on a native PAGE gel alongside reference samples that were produced separately. The resulting gel had three major bands, two of which correspond directly to bands for the self-excising module in isolation, and one of which corresponds to the RNA duplex (Figure 2e). We note that the appearance of two bands for the HHR module itself suggests the possibility of either complex formation or long-lived and distinct conformational states of the module once it has detached from the gate. From the native PAGE gel data, we conclude that the self-excising module successfully digests itself and spontaneously dissociates from the rest of the multi-stranded RNA complex.

To test if we can implement strand-displacement reactions using RNA gates generated in real-time and *in situ*, we designed a single-input single-output circuit using one self-excising module on the RNA transcript (Figure 3a). In the presence of a DNA template coding an input RNA with a toehold domain (Figure 3a, red domain), the upper part of the gate transcript is designed to be released after cleavage and subsequent toehold-mediated strand-displacement. We prepared a solution of RNA polymerase, rNTP molecules and corresponding DNA templates in a transcription buffer so that RNA transcription, folding, cleavage and displacement all happen simultaneously (Figure 3a, see Supplementary methods for more details). The output RNA was detected through recovery of the fluorescence signal of an exogenous DNA probe, via displacement of a Cy3-labelled DNA from a quencher-labelled DNA strand. When we initiated transcription using DNA templates that encode for gates with active HHRs and an input RNA, we observed a significant recovery of fluorescence within 100 minutes (Figure 3b, black solid line). In comparison, when we used the DNA template encoding gates with deactivated HHRs or removed the DNA template coding the input RNA from the reaction, no significant fluorescence recovery was observed (Figure 3b, grey solid line and black dotted line). Fluorescence recovery with the active HHR and the input RNA showed a characteristic sigmoidal behaviour: at the beginning, when not enough RNA has been transcribed yet, fluorescence recovery is slow, while after a significant amount of RNA has had the time to be transcribed, the displacement reaction occurs faster until it saturates the DNA probe.

**Figure 3.**
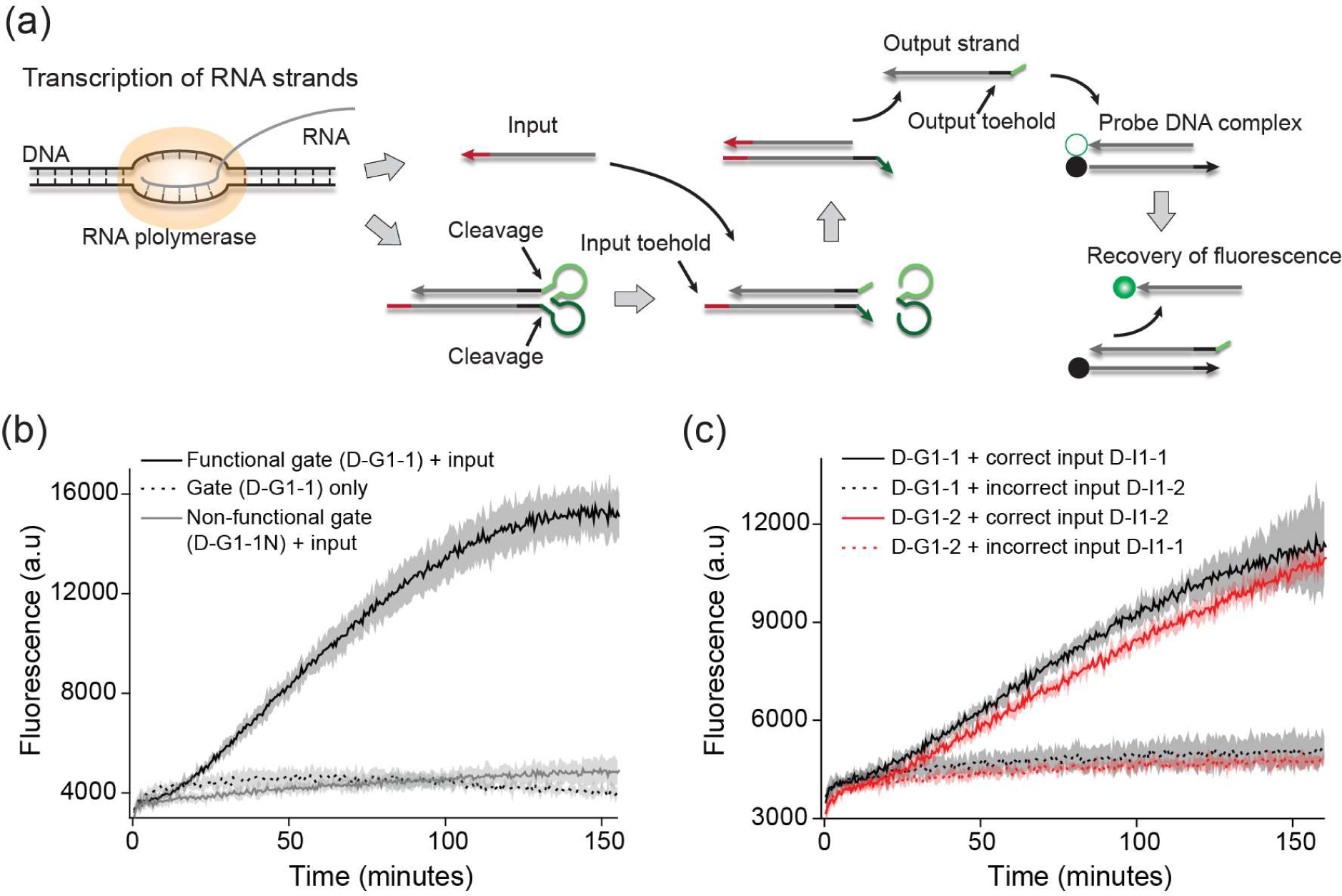
Strand-displacement reactions driven by real-time in situ enzymatic generation of RNA gate complexes. (a) RNA strands are produced continuously in the presence of RNA polymerase, rNTP molecules and their corresponding DNA templates. After folding and cleavage of the multi-stranded RNA complex, an input RNA produced simultaneously in the same volume displaces an output RNA strand from the gate complex via a toehold-mediated (red domain) strand-displacement reaction. The output RNA is detected through the increase in green fluorescence obtained when the output RNA displaces, via its output toehold (black domain), a Cy3-labelled DNA oligonucleotide from a quencher-labelled oligonucleotide. The reporter duplex is introduced exogenously unlike the components of the network itself. (b) The fluorescence increases only when the template for an input strand is present together with a functional gate. When the ribozymes are not active, or the input is absent, no significant displacement occurs. The line and shaded regions represent the mean and standard deviation from experiments in triplicate, respectively. Functional and non-functional gates are transcribed from templates D-G1-1 and D-G1-1N, respectively, and the input from template D-I1-1. (c) Orthogonality of simple strand displacement gates produced in situ. In four separate experiments, DNA templates for two gates differing only in their input toehold sequence (red domain in (a)) are transcribed alongside DNA templates for inputs possessing either correct or incorrect toeholds. The strand-displacement reactions occur only when the correct input toehold sequence is used. Solid lines correspond to templates for gates and inputs with matching toeholds (D-G1-1 with D-I1-1 and D-G1-2 with D-I1-2) while dotted lines correspond to assays with swapped input templates. The line and shaded regions represent the mean and standard deviation from experiments in triplicate, respectively. For these specific assays, 30 ng of D-G1-1 DNA template and 300 ng of D-G1-2 DNA template were used.

We tested orthogonality of our network components by exploring whether our *in situ-*generated RNA gates could discriminate between input RNAs differing only in their toehold sequences (Figure 3a, red domain). This discrimination is crucial, as many strand-displacement circuits depend on the toehold sequence to differentiate between desired and undesired reactions. Indeed, when input RNAs with interchanged toehold sequences were used, no significant recovery was observed compared to when RNAs with correct toehold sequences were used (Figure 3c, solid lines vs dotted lines). Finally, we explored the possibility of integrating *in situ* produced gates into larger reaction networks by creating and testing a two-step cascade scheme (Figure 4a). In our two-step cascade scheme, gate 1 reacts with an input strand to release an output strand that acts as an input strand for gate 2. The output of gate 2 is detected by an exogenous DNA probe. A mismatch was introduced in gates 1 and 2 and repaired by their corresponding inputs to provide thermodynamic energy drive^36^. When we tested the two-step cascade, we observed significant recovery only in the presence of the complete set of DNA templates (Figure 4b, black line vs other lines). This cascade experiment also confirms that changing the order of transcription does not prevent operation of the gates as the input toehold of gate 2 is located at the 3’ of the transcript while the input toehold of gate 1 is located at the 5’ of the transcript.

**Figure 4.**
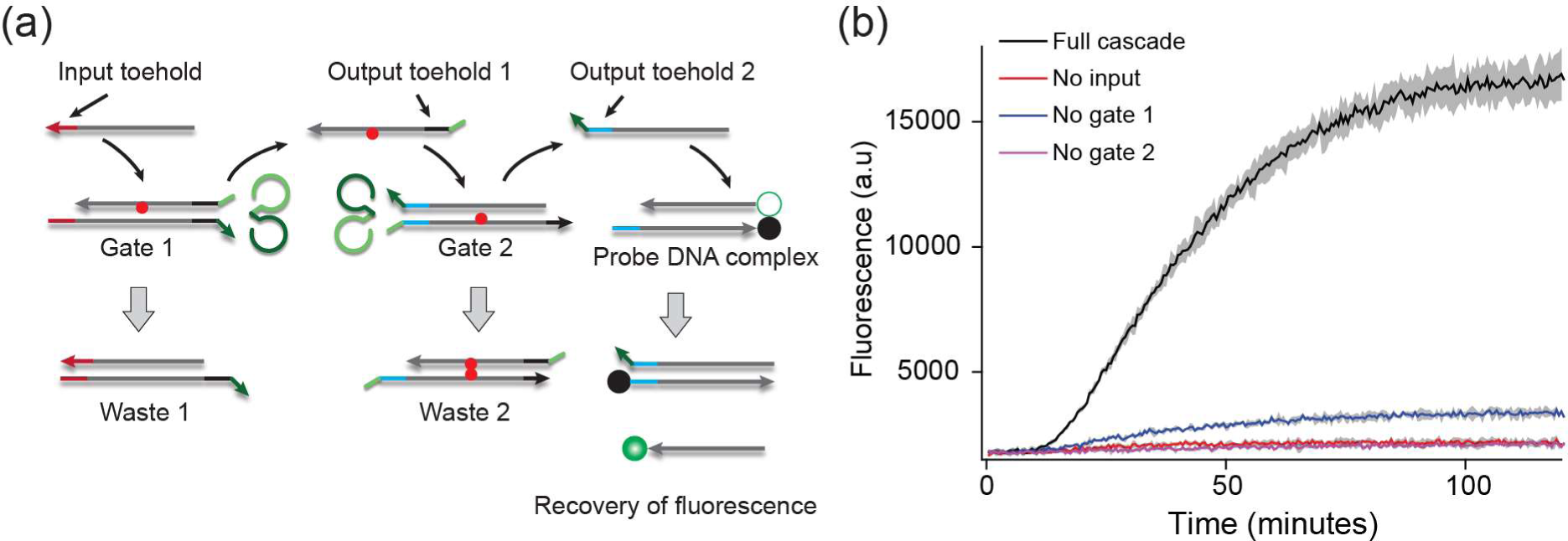
A two-step cascade system. (a) Schema of a two-step cascade system. After being triggered by an input strand using the input toehold (red domain), the output strand of gate 1 triggers gate 2 (output toehold 1, black domain) which in turn activates DNA probe with its output (output toehold 2, blue domain). A mismatch is placed on gates 1 and 2 to provide a thermodynamic drive to the circuit (red dots)^36^. All strands are produced from corresponding DNA templates (gate 1: D-I1-5, gate 2: D-G2 and input: d-I1-1) except for the probe DNA complex. (b) The fluorescence from the DNA probe increases only when all the DNA templates of the two-step cascade are present (black line compared to the other lines). The line and shaded regions represent the mean and standard deviation from experiments in triplicate, respectively.

To generalize our approach further, it is necessary to implement multi-input, multi-output gates. Doing so will require incorporating a greater number of self-excising modules on a single template and placing the module at different positions on the RNA such as in the middle of an RNA duplex. We have confirmed that the self-cleavage occurs when the self-excising module is in the middle of an RNA duplex (Figure S4) or when there are two self-excising units (Figure S5). However, the resultant gates were not fully reliable in the context of strand displacement. Indeed, it was observed that some candidate designs for our single-input, single-output gates did not function reliably (Figures 5 and S6). Further work is therefore needed to identify rules for sequence design of functional gates.

**Figure 5.**
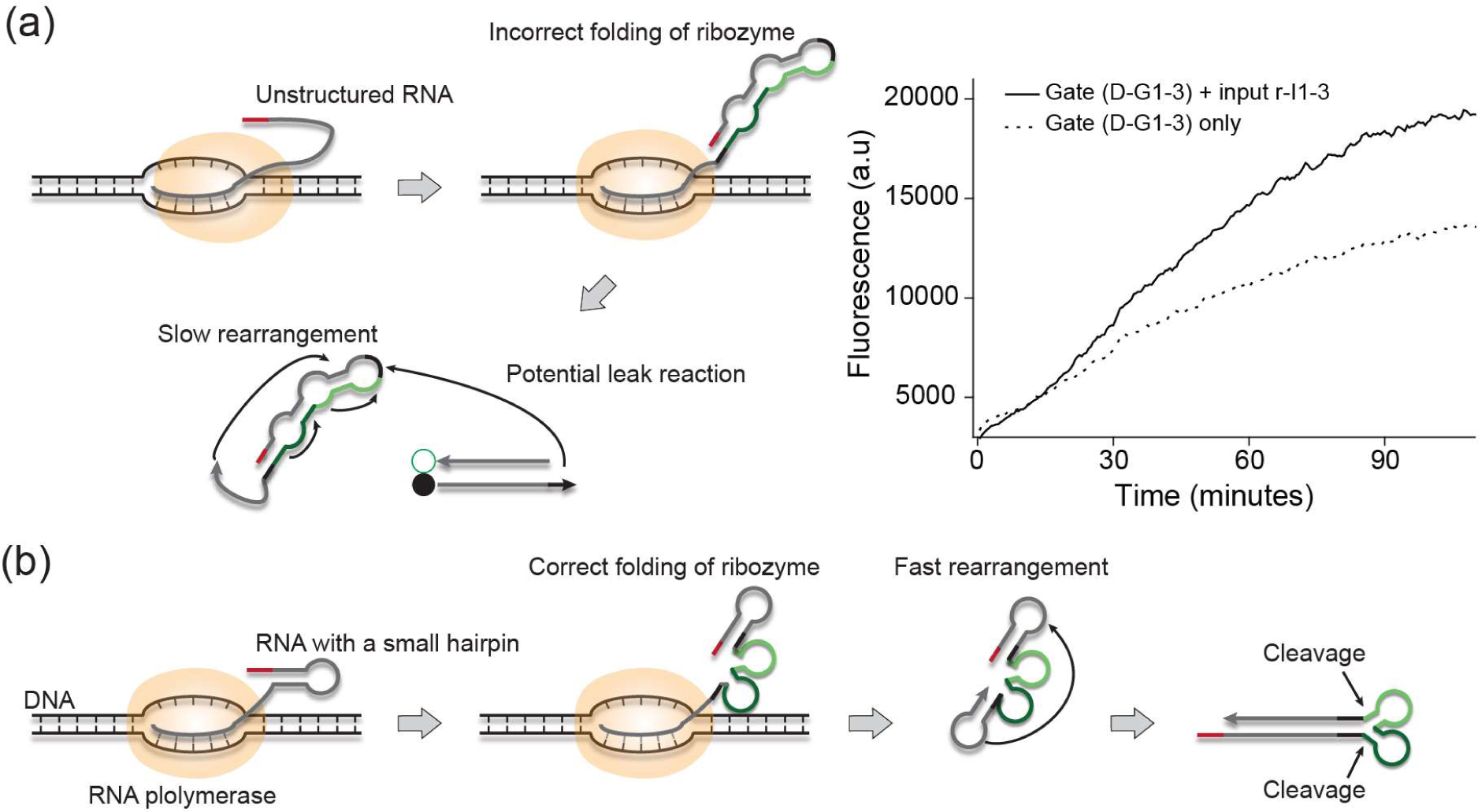
Ensuring rapid co-transcriptional folding of RNA transcript by placing a hairpin in the design of the self-excising module. Proposed mechanism to ensure rapid folding of a self-excising module and duplex domain of multi-stranded RNA during transcription. (a) If the duplex domain (grey) is unstructured, it can interact with partially transcribed ribozyme sequences and consequently form complex secondary structures that take up to minutes to rearrange, leading to potential leak reactions (graph on the right). In this experiment, gate template D-G1-3 was transcribed in solution with a fluorescent reporter and either with or without exogeneousinput RNA r-I1-3 at a concentration of 1 µM.. (b) In comparison, when a small hairpin is introduced at the 5’ end of the transcript, the ribozyme can fold correctly without interference from neighbouring sequences. Folded ribozymes should then act as nucleation sites for the complete folding of the desired structure by bringing the beginning of the duplex domain in close proximity (black domains).

PAGE gels of the malfunctioning gates, performed after 2 hours of transcription, indicate efficient cleavage (Figure S7). The issue doesn’t seem therefore to be instability of the ribozyme folding over a long timescale. We hypothesise instead that the transient kinetics of the gate folding could be a key factor in the design. During transcription, RNAs folds into its secondary structure within microseconds^37^ while the transcription of the gates typically take seconds to complete with the velocity of RNA-Polymerase elongation being around 40-50 nucleotides per second^38^. This ratio of rates suggests that the 5’ part of the RNA will form secondary structures before the 3’ part of the RNA gets transcribed, potentially trapping the transcript into a local minimum of free energy that needs to be rearranged for the RNA to fold into the global minimum of free energy that was intended by design (Figure S8). Depending on the complexity of the structure, rearrangements could take up to minutes^39^, which is slower than the speed of typical toehold-mediated displacement reactions^6^. A slow rearrangement that interferes with the operation of the ribozyme or leaves outputs of the gate exposed to interact with their downstream counterparts is potentially disastrous for the accurate implementation of circuits.

Given correct folding of the ribozymes, it is likely that the remainder of the transcript should fold quickly into the desired structure; the ribozymes effectively anchor the ends of long duplex sections in close proximity. We therefore hypothesise that it is desirable to reduce interference with the correct folding of ribozymes during transcription. The designs used in Figures 3 and 4 have a small hairpin at the 5’ end of the transcript (Figure 5). This hairpin should effectively sequester the toehold/displacement domain whilst the rest of the single-stranded RNA is being transcribed, allowing the ribozymes to fold without interference (Figure S8). When the hairpin was removed from the displacement domain by only changing the sequence of the input toehold, we observed a significant amount of leak reaction between our RNA gates and the DNA probe in the absence of the input RNA (Figure 5a). In future work, we will look into generalising this idea of directing co-transcriptional folding as we also generalise the gate motif.

In the experiments performed hitherto, reporter complexes have been introduced exogenously. In future work, we aim to produce all components *in situ*, using fluorogenic RNA aptamer systems like Broccoli^40^ as a real-time reporter. Doing so will enable us to transfer our setup to a more cell-like environment, in which components are continuously turned over by RNAse enzymes. Such a system would reach a dynamic, non-equilibrium steady state with the potential to respond to new input signals indefinitely – a more life-like setting than traditional applications of DNA nanotechnology^36^. Once these systems are optimised, it should be possible to transfer the gate motif to living cells.

## Acknowledgement

This research is supported by EPSRC grant EP/P02596X/1. TO is supported by a Royal Society University Research Fellowship. G.-B.S is supported by a Royal Academy of Engineering Chair in Emerging Technology for Engineering Biology.

## Author contributions

G.-B.S. and T.O conceived and designed research. W.B, G.-B.S and T.O designed experiments. W.B performed experiments and analysed the data. W.B, G.-B.S and T.O co-wrote the paper.

## Competing interest

The authors declare no competing interest.

## Additional information

Supplementary information is available for this paper. Raw data for the experiments can be found in the Supplementary Excel file.

## Notes

### Competing Interest Statement

The authors have declared no competing interest.

