## Supplementary material for "*In situ* generation of RNA complexes for synthetic molecular strand displacement circuits in autonomous systems": readme.txt

WB HHR SI.PDF contains supplementary methods and figures S1-S9

WB HHR rawdata.xls contains following raw data and sequence from the paper. 
-gel bands intensity for figure 2 and S3. 
-DNA and RNA sequences used in the study with colour annotation.
-Raw realtime trace data of fluorescence signal in arbitrary unit for Figure 3b, c, 4b in in three replicas and one replica for 5a, S6, S9.
