## Supplementary material for "*In situ* generation of RNA complexes for synthetic molecular strand displacement circuits in autonomous systems": WB HHR SI.pdf

### Table of contents

**Supplementary methods**

**Figure S1-S9**

### Supplementary methods

**DNA and RNA design.** All the DNA and RNA sequences including the primers that were used in this study can be found in a separate Supplementary Excel file. The DNA and RNA constructs were designed on Benchling and confirmed with NUPACK[1]. All the sequences are given in 5' to 3' order. The sequences of full-length HHRs were taken from the literature (HHR1, GGAAGAGGTCGGCACCTGACGTCGGTGTCCTGATGATGATCCATGAGAGGATCGAAACCTCTTCTAG[2], HHR2, GCAGGTACATACAGCTGATGAGTCCCAAATAGGACGAAACGCGGAAACGCGTCCTGTATTCCACTGC[3]). We used the CChMVd-U10 variant of HHR1 as it has the fastest cleavage kinetics. The double-stranded region of RNA was randomly generated except for a small hairpin domain. When designing the DNA templates, at least 4 extra DNA bases were placed upstream of the T7 promoter sequence to ensure efficient transcription of the RNA. Also, as the primers cannot bind efficiently to the template DNA during PCR if they target regions with secondary structures (for example, grey displacing domains in Figure 3a), the DNA construct was further extended at both its 3' and 5' ends. We placed a ScaI cleavage site at the 3'of the DNA sequence that encodes a double-stranded region of RNA because extra bases at the 3' affected the efficiency of the displacement reaction (Figure S9). No such cleavage was needed for 5' extensions because they are not transcribed. The primers, double-stranded DNA constructs (gBlocks), RNA and DNA oligonucleotides were ordered from IDT. All the single-stranded DNA and RNA were ordered with no additional purification except for labelled DNA strands and DNA strands to transcribe input strands which were ordered with HPLC and PAGE purification respectively.

The full sequences of the DNA templates for the gates with a colour annotation can be found in the Supplementary Excel file. To design the gate 1 series (D-G1 series), its 23 bp common displacing domain was generated first (GATTAGGAACCCAGTATCTGCAG, grey domain in figure 3a) with a 6 nt output toehold (black domain in figure 3a). An 8 nt input toehold (Figure 3a, red domain in the gate) was designed and placed at the 5' end of the displacing domain with considerations for the secondary structure (Figure S6). Then we placed the self-excising module at the 3' of the output toehold with a TT or AA spacer. We used the self-excising module with a short stem on ribozyme 1 (Figure 2b) to facilitate the detachment of the self-excising module from the gates. At this point, we used NUPACK to provide a rough check for whether the self-excising module will fold correctly. If the ribozymes failed to fold into their correct secondary structure, interfering bases in the displacing domain, output and input toeholds were mutated until we achieved correct folding as predicted by NUPACK. Then a reverse complement of the displacing domain and output toehold was added after the self-excising module with a TT or AA spacer. Finally, the promoter and spacer (TGGCAAGGTACTCACTAATACGACTCACTATAG, 5') were placed at the very beginning of the construct while the 3' reverse primer binding domain with ScaI cleavage site (AGTACTGCTAGTACGCTATTCTTTAGC, 3') was added to the very end of the construct. Sequences of input RNA for corresponding gates were determined by taking the reverse complement of the input toehold and displacing domain.

The only difference for gate 2 (D-G2-1) compared to gate 1 is the location of the input toehold which is now after the reverse complement of the displacing domain (Figure 4a, black domain). The 6 nt output toehold (Figure 4a, blue domain) was designed and added after the displacing domain with similar consideration for the secondary structure above. After adding the self-excising unit with an AA spacer after the output toehold, folding of the partial structure consisting of the displacing domain, output toehold and self-excising unit was checked with NUPACK. The reverse complement of the displacing domain and output toehold was placed after the self-excising unit with an AA spacer. Then the input toehold (output toehold from gate 1, Figure 4a, black domain) was added after the displacing domain. Finally, the same promoter with a spacer and the 3' primer binding domain used above were added at the 5' and 3' of the construct respectively. To provide a thermodynamic drive to the reaction, a mismatch, which gets repaired by the input RNA strands, was introduced in the displacing domain of the gates (Figure 4a, red dots).

The DNA templates to transcribe the input and longer RNA fragments (Figure 2e, 3 and 4) were designed by adding a T7 promoter (TAATACGACTCACTATAG) with a 4 bp spacer at the 5' upstream of the required sequence. As the input DNA templates were too short for gBlock synthesis, their forward and reverse strands were ordered as long single-stranded DNA oligonucleotides from IDT with PAGE purification. Then the oligonucleotides were thermally annealed at 50 µl concentration by heating at 90°C for 1 minute and cooling down to room temperature over 1 hour. Probe DNA Cy3-quencher (Iowa black FQ) pairs, designed to bind the output toehold and displacing domain (Figure 3a, black and grey domains and Figure 4a, blue and grey domains) were ordered with HPLC purification from IDT and then annealed using the same protocol.

**DNA assembly and amplification.** DNA constructs with a small degree of secondary structure (D-H1, D-H2, D-SEU1 and D-SEU2 series) were synthesized directly using a single gBlock and amplified as described later. DNA constructs with longer double-stranded regions or long repeats of RNA (D-G1, D-G2 series and D-SEU12) could not be synthesized from single gBlock synthesis. Such constructs were split at a location that minimized complexities with overhangs for the Golden Gate assembly scheme. Specifically, the overhangs were designed for the BsaI enzyme restriction site with at least 4 bp spacers to ensure high yield of digestion. PCR amplification of each gBlock was done with Q5 DNA polymerase (NEB) at the annealing temperature specified in the Supplementary Excel file. After purification using the Monarch® PCR Purification Kit (NEB), the sub-components were incubated at 1:1 molar ratio in the presence of BsaI (BsaI-HFv2 from NEB) and T7 DNA ligase (NEB) for 10 cycles of digestion – 5 minutes at 37°C – and ligation – 30 minutes at 16°C in a thermocycler (Thermofisher). The assembled DNA was run on a 2% native agarose gel and purified using the Monarch® DNA Gel Extraction Kit (NEB). The purified DNA was used as a template for another round of PCR amplification and then purified as necessary. The DNA fragments were PCR-amplified with a higher concentration of primers (1.5 μM each) and smaller number of cycles (typically less than 15 cycles) to minimize secondary bands that appear at higher molecular weight. To detect such unwanted band with high resolution, the amplified DNAs were run on a pre-cast 10% native PAGE gel system from Invitrogen (Novex gel system) at 200V for 30 minutes with SYBR safe staining (Invitrogen). If a purification step was required, the band of interest were sliced from the gel and eluted by crushing and soaking in 5 times excess of TAE buffer. The DNA was then concentrated with a PCR purification/concentration kit (Monarch® kit from NEB). If only a correct band was observed, the PCR samples were directly purified with PCR purification/concentration kit to maximize the yield. After preparation of the DNA template, the extra 3' domain was cleaved with ScalI if necessary. The concentration of purified DNA was measured by a Nanodrop (Thermo scientific).

**RNA transcription and characterization.** All RNA transcription assays used the following protocol, adapted from the standard protocol for *in vitro* transcription using T7 RNA polymerase from NEB unless stated explicitly in the figure caption. The RNA was transcribed from the DNA templates using T7 RNA polymerase from NEB using standard protocols with a slight modification detailed hereafter. For ribozyme characterization assays, 100 ng of the template DNA was added for each 20 μl of transcription reaction. To maximize the amount of RNA transcribed, we increased the concentration of rNTP monomers from 0.5 mM to 1 mM and supplemented the RNA transcription buffer (NEB) with 5 mM of $MgCl_2$. After 2 hours of transcription at 37°C, the template DNA was digested by adding DNAse 1 (NEB). The transcribed RNA was then purified using the GeneJet RNA concentration kit (Invitrogen) and quantified with a Nanodrop (Thermofisher). The typical yield was 6 ng of RNA per 20 μl of transcription reaction. Then 1 μg of the purified RNA samples were denatured by incubating for 10 minutes at 70°C after adding 2X RNA gel loading dye (Thermofisher). The denatured samples were run on a precast 15% TBE-urea PAGE gel (Novex gel system, Invitrogen) for 30 minutes at 200 V alongside with 1.5 μl of the low molecular weight marker (NEB). The gel was stained with single-stranded specific staining dye SYBR green 2 (Invitrogen) for visualization. The image was analysed with ImageJ. The intensity of the bands and the dark background were measured separately using the measurement function of ImageJ. After subtraction of the dark background, the yield Y was estimated by $Y = \frac{cut}{uncut+cu}$. Raw intensities and estimated yields can be found in the Supplementary Excel file. For the gate dissociation test, the RNA samples were loaded (Gel loading solution, Invitrogen) and run in 15% TBE PAGE gel (Novex gel system, Invitrogen) at room temperature for 30 minutes at 200 V. In the dissociation test, control RNA 1 and 3 were synthesized from IDT while control RNA 2 was transcribed from the corresponding DNA template from IDT.

**Strand displacement reactions using complexes produced *in situ*.** All the strand-displacement assays were performed using the following protocol, except where explicitly stated in a figure caption. In a 96-well plate (Greiner, black and μClear plate), we prepared a 50 μl reaction solution containing 1X of T7 RNA polymerase and transcription buffer (NEB), 0.5 mM of each rNTP monomers (NEB), 50 ng of template DNA for gate, 50 ng of template DNA for input RNA and 50 nM of Cy3-quencher probe pair. The fluorescence was monitored by a microplate reader (BMG, Clariostar) using 530 nm excitation and 580 nm emission wavelength every 30 seconds. The reaction was performed at 42°C to minimize the rearrangement time of RNA secondary structures during transcription and folding which was hypothesised to lead to leak reactions. For the two-step cascade experiment, 50 ng of template DNA for gate 1and gate 2 were added to the reaction.

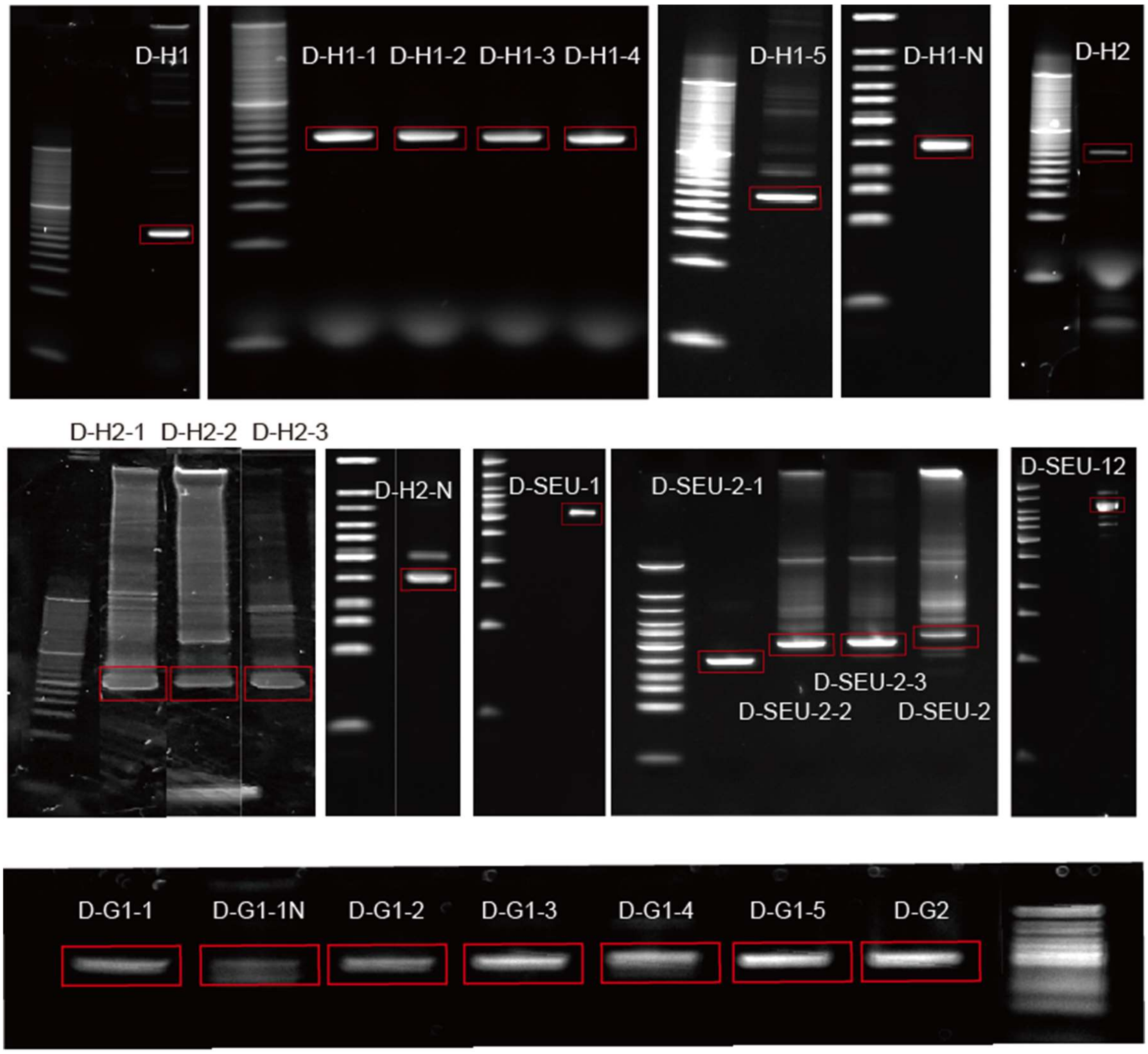


**Figure S1. Preparation of DNA templates for *in vitro* transcription.** After assembly and PCR amplification DNA templates were run on a 10% PAGE gel. For the constructs with clear single band, the PCR product was purified directly with a PCR clean up kit (Monarch, NEB). For the constructs with more than one band (D-H2 series, D-SEU series), the band was excised from the PAGE gel and the DNA inside was purified by soaking overnight in 1X TE buffer with five times the volume of the gel. The DNA in the TE buffer was concentrated using a PCR purification kit. Full name and description of each DNA template can be found in the Supplementary Excel file.

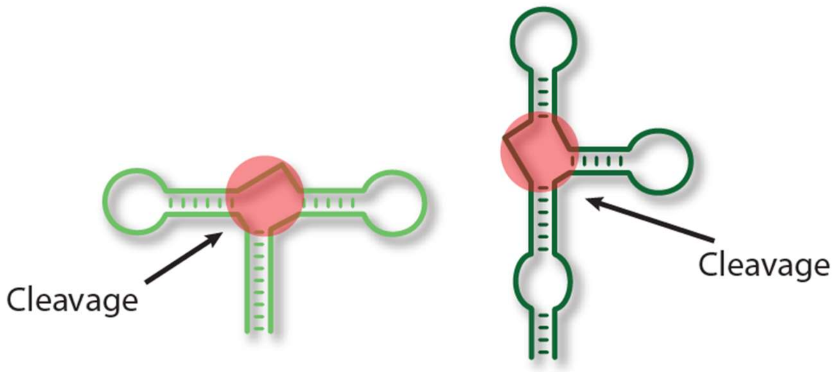


**Figure S2. Non-functional version of HHR1 and HHR2.** The cleavage activity of each HHR was inactivated by replacing sequences in the catalytic core of each HHR (red regions). Specifically, all the purines were changed to pyrimidine and *vice versa*. The full sequence can be found in the Supplementary Excel file.

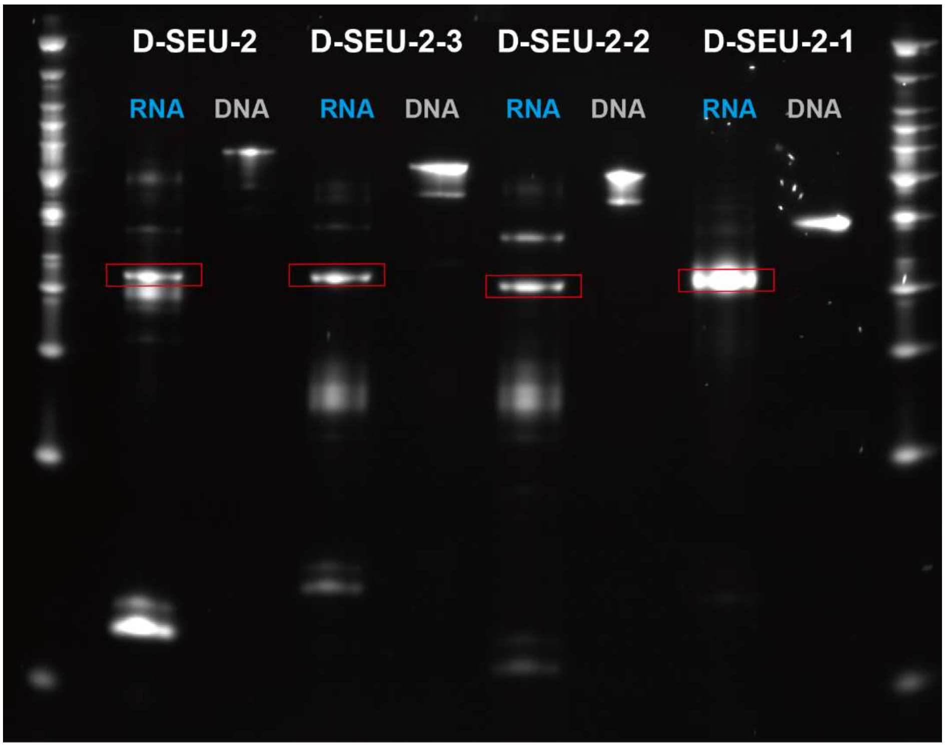


**Figure S3. Testing the cleavage yield of the engineered self-excising module.** The RNA transcript from the engineered self-excising module designed with randomized or shorter stems were run alongside its DNA template. The boxed bands are consistent with the intended size of the self-excising unit (108 nt), indicating successful cleavage. Randomizing the sequence of each stem (D-SEU-1, D-SEU-3) or shortening the length of one stem (D-SEU-2) does not hamper cleavage significantly relative to the default D-SEU-2.

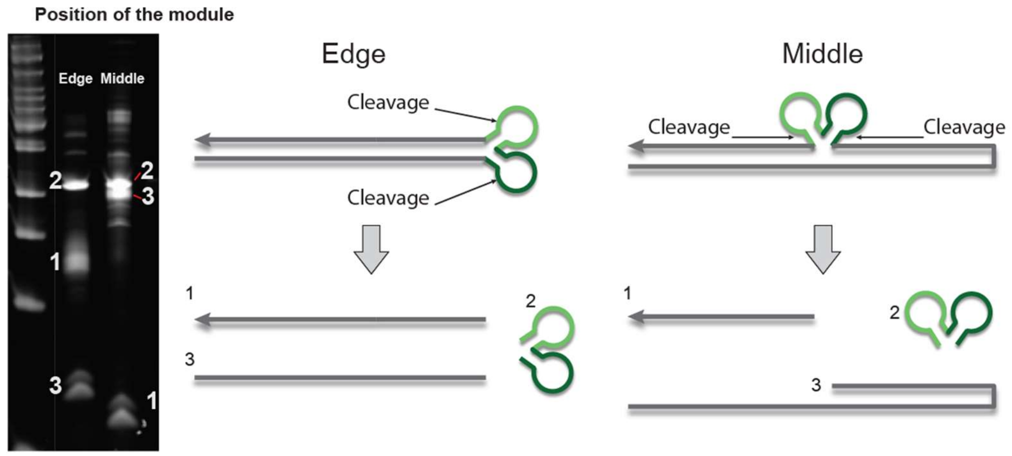


**Figure S4. Successful cleavage of the self-excising unit when placed in the middle of an RNA duplex.** Three distinct bands appear in a denaturing PAGE gel when the self-excising unit is placed on the edge (D-SEU1) or in the middle (D-SEU2) of an RNA duplex. These bands appear approximately where they are expected.

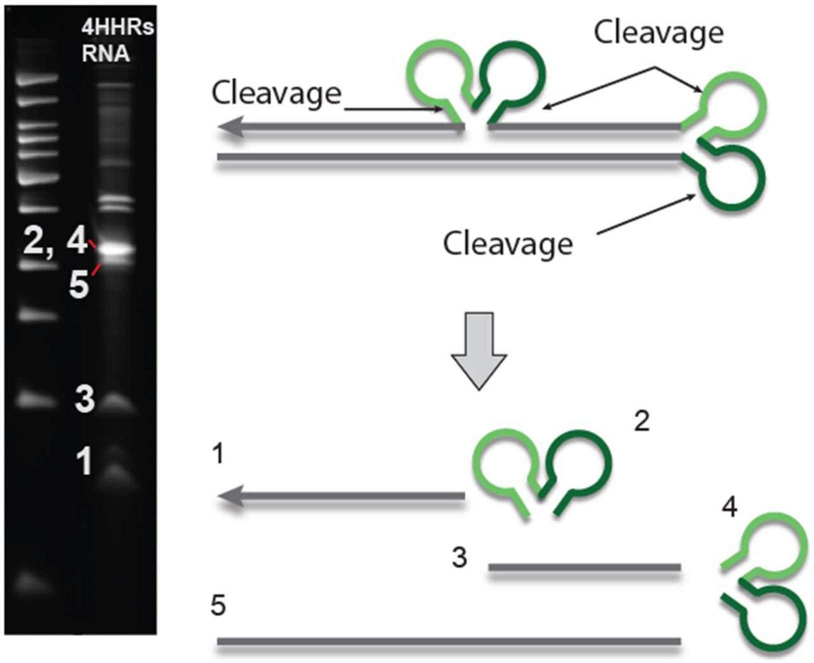


**Figure S5. Successful cleavage of the gates with two self-excising units (D-SEU12).** 4 distinct bands appear in a denaturing PAGE gel when the self-excising unit is placed in the middle of an RNA duplex. The self-excising modules appear in the same position.

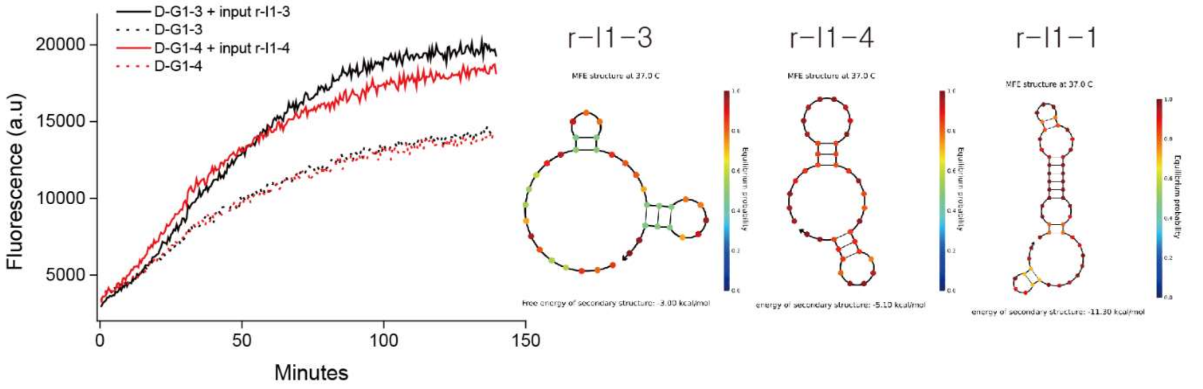


**Figure S6. Strand-displacement reactions driven by *in situ* enzymatic generation of RNA gate complexes with no hairpin in the displacement domain function poorly.** We observed significant leak from the gates in the absence of their input RNA (left graph) when a hairpin is removed from the secondary structure of the gates. The secondary structures of the first 37 nt of the gates which contains its input toehold, displacing domain and output domain are drawn on the right. The correctly functioning gate template (D-G1-1) has a stronger hairpin in its RNA transcript than the poorly-functioning gate templates (D-G1-3, D-G1-4). For the input-triggered reactions, 1 μM of exogeneous RNA input (r-I1-3 or r-I1-4) was added to the template and reporter transcription mix instead of the corresponding DNA template..

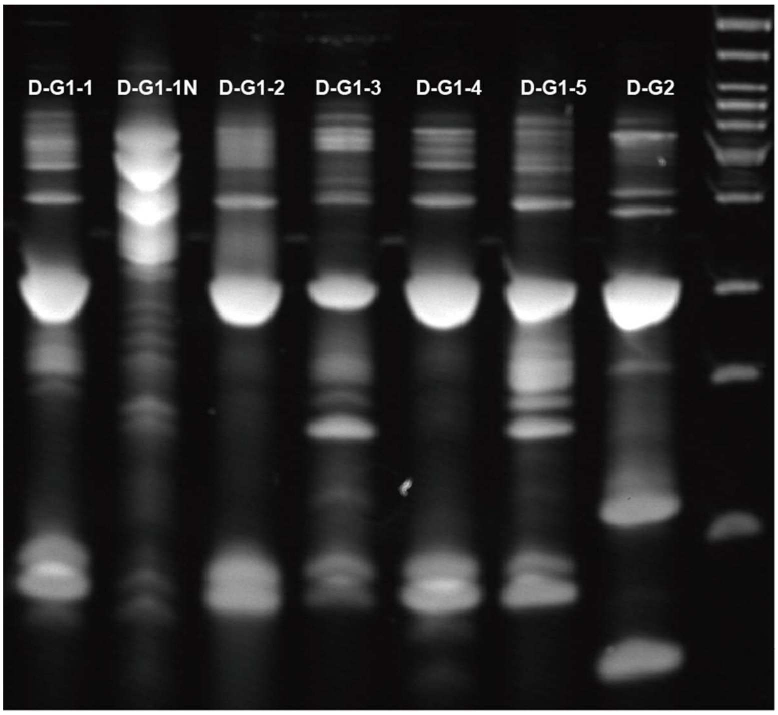


**Figure S7. Successful cleavage of self-excising modules in gates used for the *in situ* experiments.** After two hours of transcription, 5 μl of each transcript was run on the 15% TBE-Urea PAGE gel. Successful cleavage are observed compared to non-functional variant of the gate 1 (D-G1-1N). Full sequences and description of each DNA template can be found in the Supplementary Excel file.

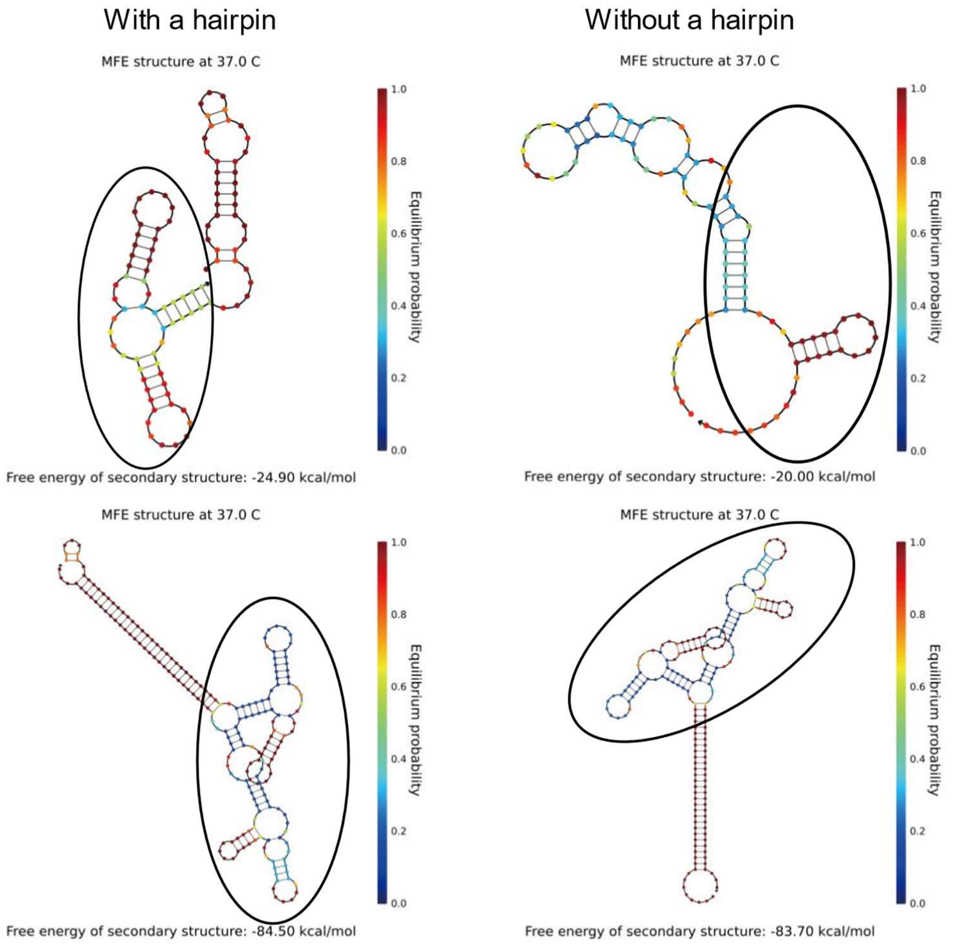


**Figure S8. Secondary structures of partial and complete RNA transcripts, with and without small hairpins in the duplex domain.** NUPACK[1] was used to predict the secondary structures of the first 97 nt with a single-stranded displacing domain and one hammerhead ribozyme (upper images) as well as full length RNA transcripts (lower images). Only when a small hairpin is introduced in the displacing domain (D-G1-1, top left, black ellipsoid), is the correctly-folded hammerhead ribozyme present in the lowest free-energy structure during the initial phase of transcription. In the absence of a hairpin, the displacing domain interferes with the folding of the HHR (D-G1-3, top right, black ellipse). Secondary structures of full RNA transcripts contain correctly folded HHRs (lower images, black ellipse). Whilst these free-energy calculations at the secondary structure level are limited in their ability to describe the kinetic behaviour of a transcript with complex tertiary structure, they are nonetheless suggestive of problematic folding of the ribozyme in the hairpin-free case.

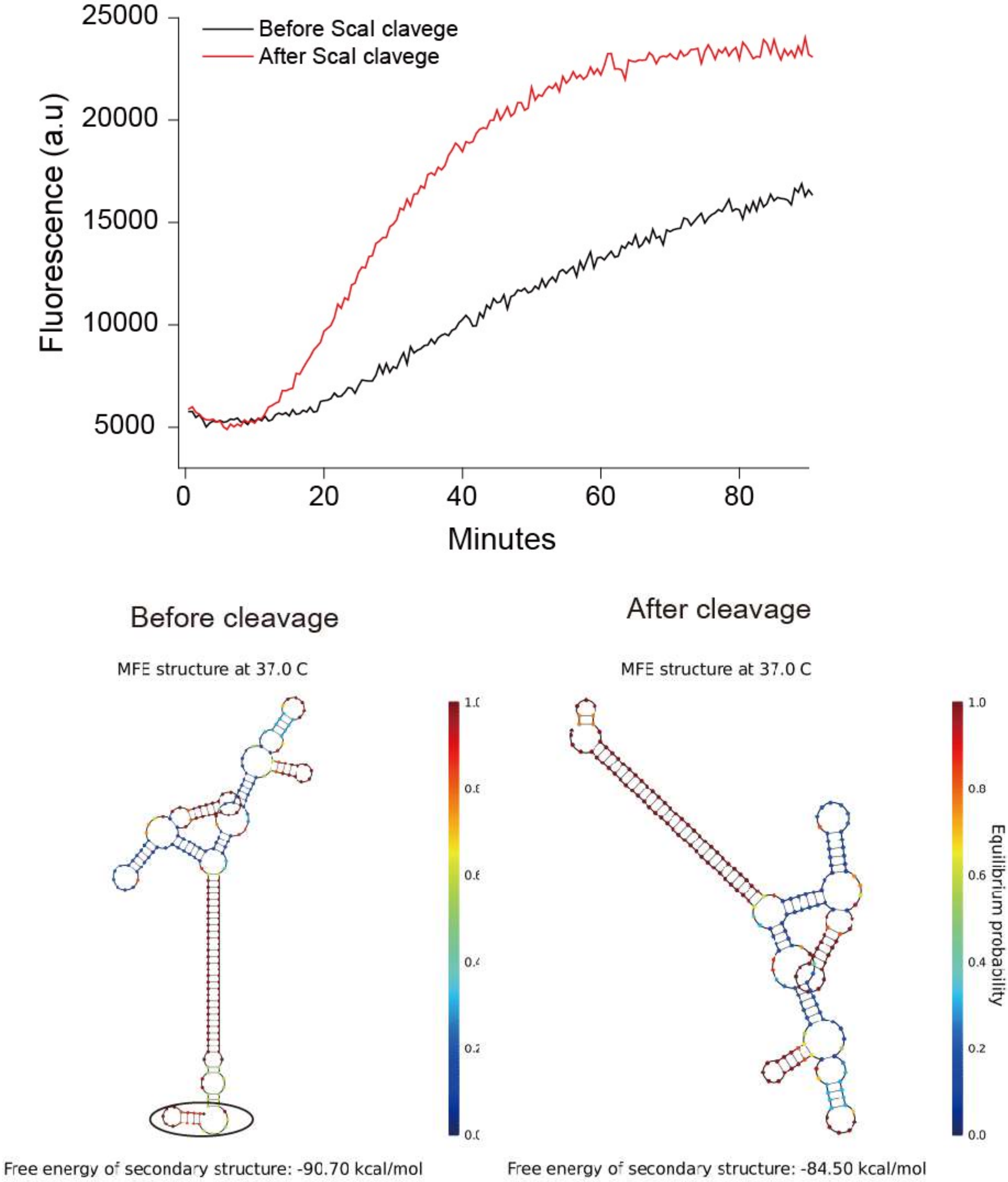


**Figure S9. Effect of cleaving the 3' extra domain of a gate during an *in-situ* strand-displacement reaction.** When the extra 3' domain (black circle) was cleaved from D-G1-1 with ScaI, the strand-displacement reaction became faster and more efficient. 1 µM of input RNA was added instead of the corresponding DNA template as an input.


1. Zadeh, J. N.; Steenberg, C. D.; Bois, J. S.; Wolfe, B. R.; Pierce, M. B.; Khan, A. R.; Dirks, R. M.; Pierce, N. A. *J Comput Chem* **2011,** 32, (1), 170-173.
2. De la Peña, M.; Gago, S.; Flores, R. *The EMBO Journal* **2003,** 22, (20), 5561-5570.
3. Osborne, E. M.; Schaak, J. E.; Derose, V. J. *Rna* **2005,** 11, (2), 187-196.
